# Neuronal networks quantified as vector fields

**DOI:** 10.1101/2024.06.29.601314

**Authors:** Szilvia Szeier, Henrik Jörntell

## Abstract

The function of the brain function is defined by the interactions between its neurons. But these neurons exist in tremendous numbers, are continuously active and densely interconnected. Thereby they form one of the most complex dynamical systems known and there is a lack of approaches to characterize the functional properties of such biological neuronal networks. Here we introduce an approach to describe these functional properties by using its activity-defining constituents, the weights of the synaptic connections and the current activity of its neurons. We show how a high-dimensional vector field, which describes how the activity distribution across the neuron population is impacted at each instant of time, naturally emerges from these constituents. We show why a mixture of excitatory and inhibitory neurons and a diversity of synaptic weights are critical to obtain a network vector field with a structural richness. We argue that this structural richness is the foundation of activity diversity in the brain and thereby an underpinning of the behavioral flexibility and adaptability that characterizes biological creatures.

## 1 Introduction

Understanding the collaboration between multiple neurons remains a challenging issue within neuroscience. Even though cortical activity is high-dimensional, standard approaches typically rely on performing some type of dimensionality reduction on the neuron population activity data [1, 2]. Dimensionality reduction results in a loss of the underlying information present in the brain circuitry signals, which could be critical to understand the neuronal interactions of the brain circuitry processing [3]. Jazayeri and Ostojic [1] compared different dimensionality reduction methods, which could be expected to report different results from the same underlying data, and arrived at the conclusion that, ‘without concrete computational hypotheses, it could be extremely challenging to interpret measures of dimensionality’. Put in other words, if we don’t know the computational architecture of the brain circuitry, it is not possible to get a dimensionality reduction right, or ‘loss-less’, regardless of the embedding method or dimensionality reduction tool used. Hence, there is a need for a conceptual framework to quantify neuron population level interactions, while not discarding the bulk of the information due to arbitrary impacts of the particular frame of reference (‘embedding’) used to approximate the nature of those interactions.

In addition to being high-dimensional, most networks in the brain are recursively interconnected, as in the neocortex [4, 5, 6, 7, 8, 9, 10] and in the spinal cord inside its circuitry as well as via its sensorimotor feedback loops [11]. The specific function of any given neuron depends on the current activity of other neurons, which means that its function will vary and be context– or state-dependent. Therefore, network operation can be nearly impossible to understand by observing activity neuron-by-neuron. It would then be more informative to think of the neuron population activity as residing in a state space defined by the combined activity of the population, and that each individual neuron is contributing to the current location within that state space through its synaptic interactions with other neurons. The location within the network state space may appear an abstract term, but in fact it is critical for the brain in defining how the spatiotemporal patterns of neuron activity will evolve in the network, thus for example directly translating to understanding how sequences of muscle activation patterns in complex movements are being formed [12, 13].

Here we wanted to focus on identifying a ‘ground truth’ conceptual framework within which such network-global multi-neuronal interactions could be understood as an emerging property of the fact that synaptically connected neurons will inevitably impact each others activity levels. The idea of this conceptual framework is that it can serve as a basis for new tools to characterize and understand multi-neuron interactions. We introduce the method of describing the activity– and weight-dependent neuronal interactions using vector field representations. We show how this representation can be used to understand the factors that impact the structure of the state space of the neuronal network, regardless of its dimensionality (number of neurons). We show how the distribution of the synaptic weights across the network shape its vector field and that inhibitory neurons are important to achieve a diversity of operational states within the network state space. We also demonstrate how subspaces, specific planes in the high-dimensional vector field, can be extracted for visualization purposes and to explore in greater detail factors that impact the multi-neuronal interactions in a particular context, such as in a specific movement phase.

## 2 Methods

### 2.1 Non-spiking neuron model

Transmission between neurons in a biological network is possible through synaptic connections. To construct our networks, we used a previously published non-spiking neuron model [14], which is an emulation of a conductance-level neuron model with a static leak component representing the leak channels of the neuron membrane (1A). The original model also has a membrane time constant (’dynamic leak’), which is omitted here where we were only interested in modelling the impact of the synaptic activity in the static setting. The activity of the post-synaptic neuron depends on the activities of the presynaptic neuron(s) and can be formulated as

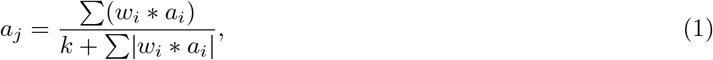

where *a*_*j*_ is the activity of the postsynaptic neuron, *a*_*i*_ represents a presynaptic neuron activity, *w*_*i*_ is its synaptic weight, and *k* is the static leak constant. In all experiments (unless stated otherwise), *k* was set to 0.1 * (*n* − 1), where *n* is the number of neurons in the network.

### 2.2 Neural state space

The impact exerted by one (presynaptic) neuron on another (postsynaptic) neuron is proportional to the presynaptic activity and synapse weight. Synaptic weights were set to [0 … 1] for excitatory synapses and to [−1 … 0] for inhibitory neurons; autapses and parallel synapses were not allowed. A weight of 0 can be interpreted as a silent synapse or the absence of synaptic connectivity. The activity of individual neurons is bounded between 0 (no activity) and 1 (saturation or epilepsy). We can define a (bounded) state space where each dimension represents an individual neurons activity level. Such a state space can be constructed for a network of arbitrary dimensionality. Essentially, we obtain a hypercube with the dimensionality being equal to the number of neurons in the network. Every state (point) in the state space, represents a neural activity distribution, that is, uniquely determines the activity configuration of all the neurons in the network. When considering short time intervals (or a single time point), the synaptic weights can be considered static.

### 2.3 Two-dimensional planar subspace

We can consider two-dimensional, planar subspaces (or extracted plane) of the original, high-dimensional neural state space. The dimensions of these subspaces represent the activity of neurons: either two individual neurons at a time or a population of neurons divided into two groups, one per dimension. When the axes represent individual neurons, the extracted plane reflects the relationship between those two neurons. However, a dimension general to representing some linear combination of neurons. Therefore, the extracted plane shows the relationship between the two selected linear combinations of neurons in the network.

## 3 Results

### 3.1 Vector field representation and plane extraction

We started out with a setting of synaptically connected excitatory cortical neurons (Figure 1A) to introduce the concept of how neuronal interactions can be quantified as vector fields. As indicated in Figure 1A we used a non-spiking neuron model with identical, linear-like neuronal input-output functions (emulating the conductance-level biological neuron behavior [14]) across all neurons. Figure 1B illustrates a network with only two reciprocally connected excitatory neurons. If a neuron is connected to another neuron with a synapse, then that means that the activity of the second neuron is dependent on the activity of the first neuron, as shown in Figure 1C at y=0 along the x-axis. Hence, the first neuron will increase the activity of the second neuron, if the synapse is excitatory. That impact will depend on the activity level of the first neuron – if that activity is very low or zero, then it will have no or very low impact on the second neuron, and vice versa for high activity. It should be noted that with the neuron model we employed, the series of vectors along the x-axis at y=0 increased at a decelerating pace because at high input activity the conductances to some degree short-circuit the impact of additional inputs [14] (Figure 1C).

**Figure 1:**
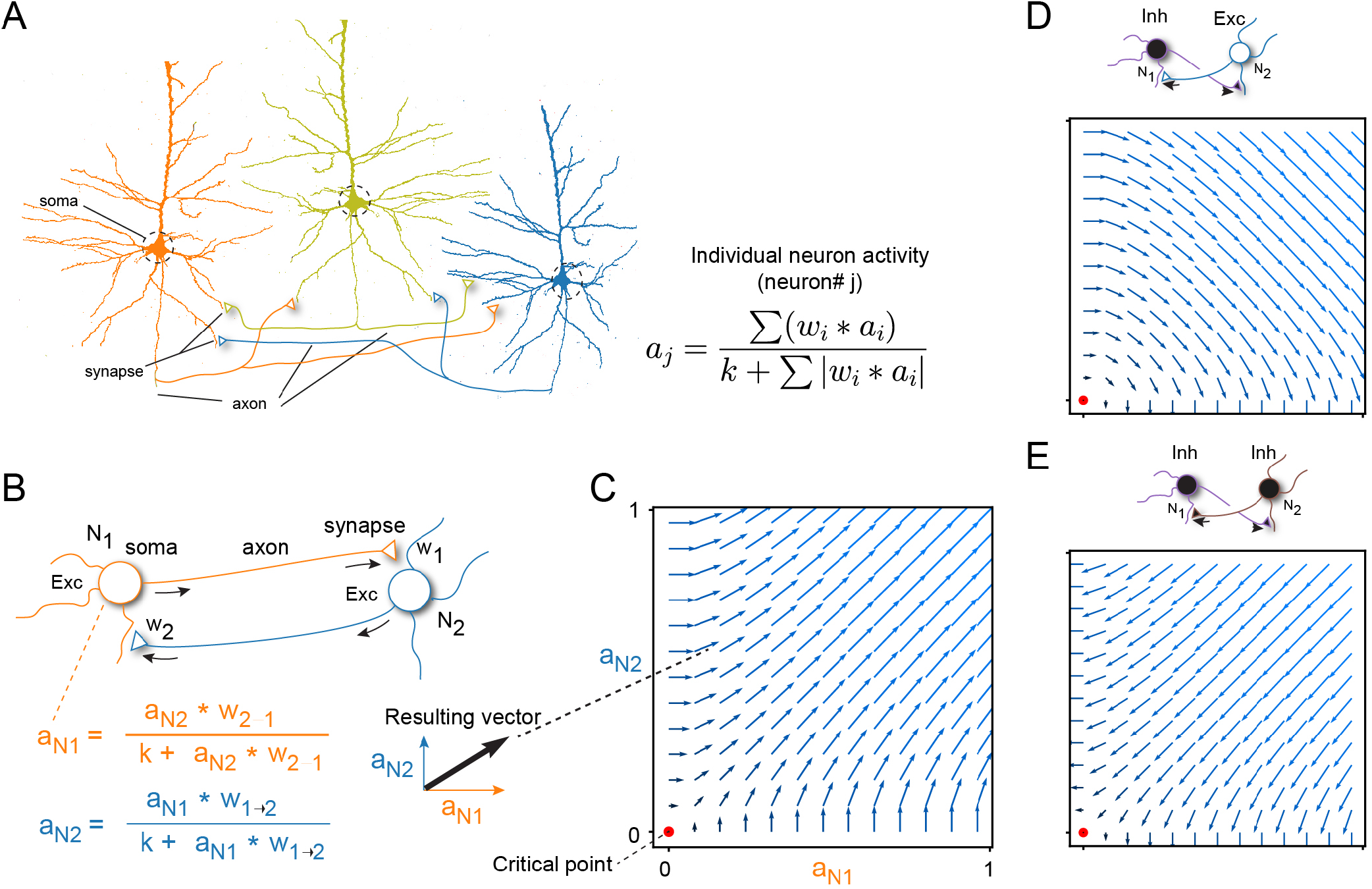
How neuronal networks map to vector field representations. (A) Actual neurons have dendrites, a soma, and an axon. The axon branch profusely and make synapses on other neurons. The example shows outlines of three (excitatory) pyramidal neurons (dendrites and axons are truncated) from previous histological analysis, as well as putative synaptic connections between the three. The inset shows the neuron model used, i.e. how the activity of each neuron was calculated. (B) Representation of the network formed between two of the pyramidal neurons. Arrows indicate the flow of signals. At each activity level, we calculated a vector from the activity levels of the two neurons as indicated. (C) Resulting vector field representation of the the two-neuron network. Each axis represents the activity of one of the neurons. (D) Vector field representations of an excitatory-inhibitory network. (E) Same as D but for an inhibitory-inhibitory network. Across all vector fields, synaptic weights were uniformly set to 1 (negative sign for connections originating from inhibitory neurons) and *k* was set to 0.2

At the same time, the activity of the second neuron will also impact the activity of the first neuron in a similar manner, as they are reciprocally connected. The interaction between the two neurons creates a vector component indicating with what magnitude the two neurons impact each other, at any possible activity level (Figure 1B). This process can then be iterated for all activity combinations across the two neurons, each combination resulting in a vector component. The resulting vector field across all possible activity combinations for the two neurons is shown in Figure 1C. As the two neurons form a positive feedback loop, they will tend to push each others activities up towards the upper right corner, i.e. this would correspond to saturation or overexcitation of the neuronal activity [15] (which in our neuron model occurs at an activity level of 1, corresponding to a maximum spiking probability of 1, as illustrated for example in [16]). In cases where one or both neurons are instead inhibitory (Figure 1D,E), the structure of the vector field will reorganize accordingly. When both neurons are inhibitory (Figure 1E), the vectors point downwards relative to the two axes and would hence tend to drive the activity of both neurons towards zero. Notably, the only neutral position within any of these vector fields, i.e. the position where the vector lengths approach zero, is when both neurons have zero activity (‘critical point’)(Figure 1C-E).

When the number of neurons connected to each other increases, this means that the dimensionality of the network increases. First, consider a network of three excitatory neurons (Figure 2A). Each neuron will be impacted by the synaptic input activities of the other two neurons. Using the same approach as above (Figure 1C), we can calculate the now 3-dimensional vector field (Figure 2B). However, it is harder to visualize the vector field in 3D, and if we add more neurons to the network it becomes impossible. For visualization purposes, we can extract a two-dimensional subspace (i.e. a plane) from the full 3-dimensional network space to visualize the interactions between two selected neurons (Figure 2C) which will now look analogous to the 2-dimensional networks of Figure 1. Similarly to the 3-dimensional case, we can extract planes from a network consisting of 4 neurons (Figure 2D-E) and from 8-dimensional networks (Figure 2F-G). This concept of plane extraction can be generalized to networks of arbitrary dimension.

**Figure 2:**
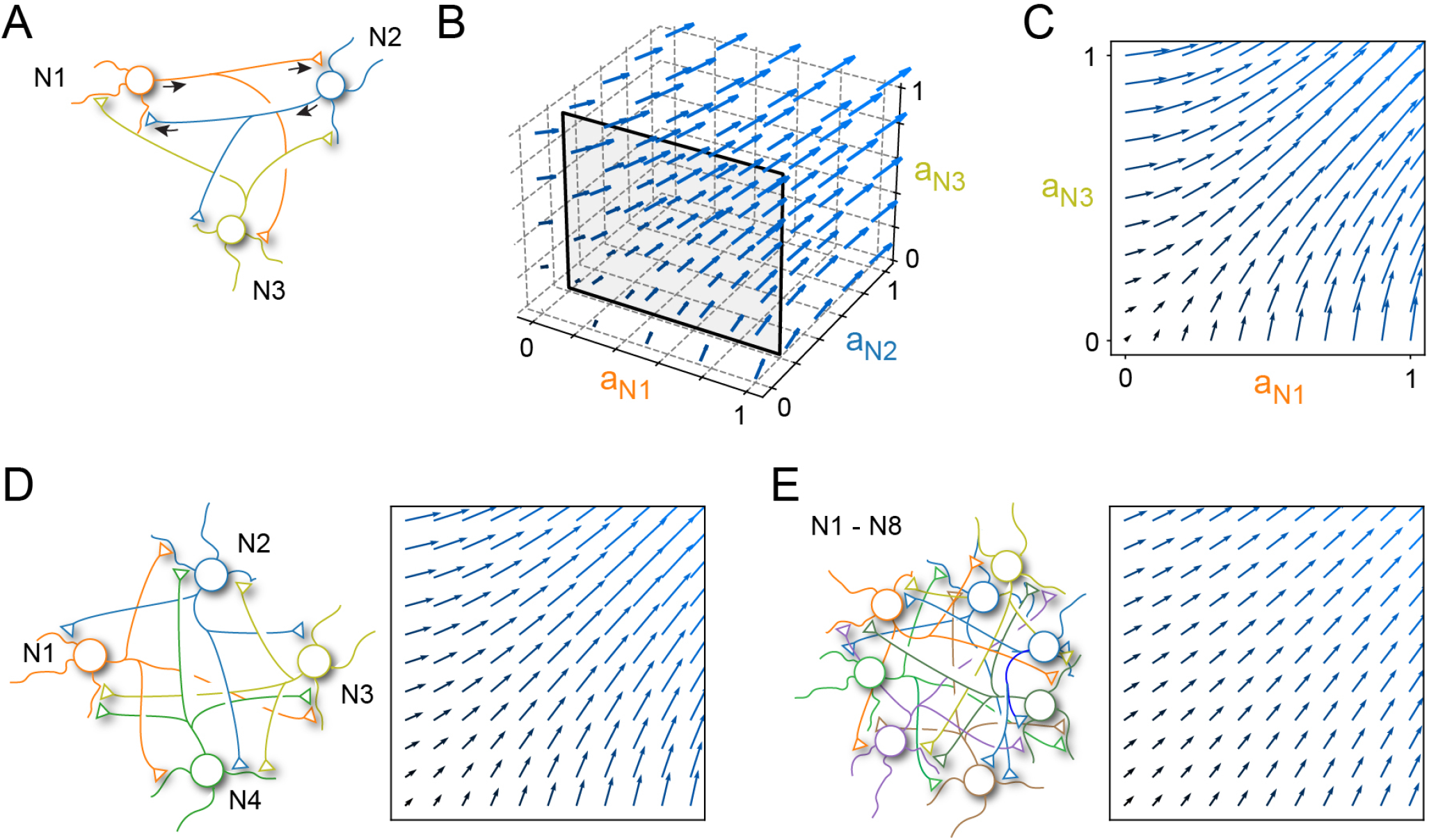
Extending the vector field representation to higher dimensions. (A) Representation of the excitatory network formed between the three pyramidal neurons. (B) Vector field representation of the three-neuron network. (C) A 2D plane extracted from the 3D vector field, corresponding to the plane where the activity of the perpendicular neuron (N2) was fixed. (D) Network consisting of four pyramidal neurons and an example plane extracted vector field from the 4-dimensional vector field space. (F) An eight pyramidal neuron network with an example extracted plane. Across all panels, the synaptic weights were uniformly set to 0.1. The perpendicular neurons were set to have an activity of 0.1.

When we choose to visualize a plane, that is equal to fixating the activity of the neuron that is not represented on the two axes of the plane. We refer to this neuron as the ‘perpendicular neuron’ since its activity axis is perpendicular to the axes of the visualized plane. If we vary the activity level of the perpendicular neuron, it will lead to different vector fields in the visualized plane, as the influence of the perpendicular neuron on the other neurons will change with its activity level. Note that the critical point could end up being be located outside the visualized plane, if any of the perpendicular neurons had non-zero activity, which indeed was the case for all planes illustrated in Figure 2.

The two dimensions of the visualized plane can also represent a linear combination of neurons in the network, rather than being limited to a single neuron per axis. In the same 3-neuron excitatory network as before (Figure 3A), we can orient the plane in its 3-dimensional vector field such that one of its axes represents the combined activities of two out of the three neurons, i.e. the axis is orthogonal to only one of the neurons (N3 in Figure 3B). The x-axis of the extracted plane now represents a weighted linear combination of the first two neurons (Figure 3C), with each neuron’s contribution determined by the angle of the plane’s orientation relative to the neuron activity axes. Since this axis now corresponds to the combined activity of multiple neurons, its maximum value can exceed 1, even though the activity of any single neuron cannot (Figure 3C, red arrows). This concept of plane extraction can be extended to networks of arbitrary dimension (Figure 3D), where each of the two dimensions (axes) of the plane can represent any combination of neurons (Figure 3E,F). In fact, a single neuron could even be represented on both dimensions of the extracted plane, if both the axes of the plane are oriented at non-perpendicular angles to the activity axis of that neuron. This plane visualization can be used to examine in detail the impact that a network parameter change, such as a specific synaptic weight, would have on the vector field.

**Figure 3:**
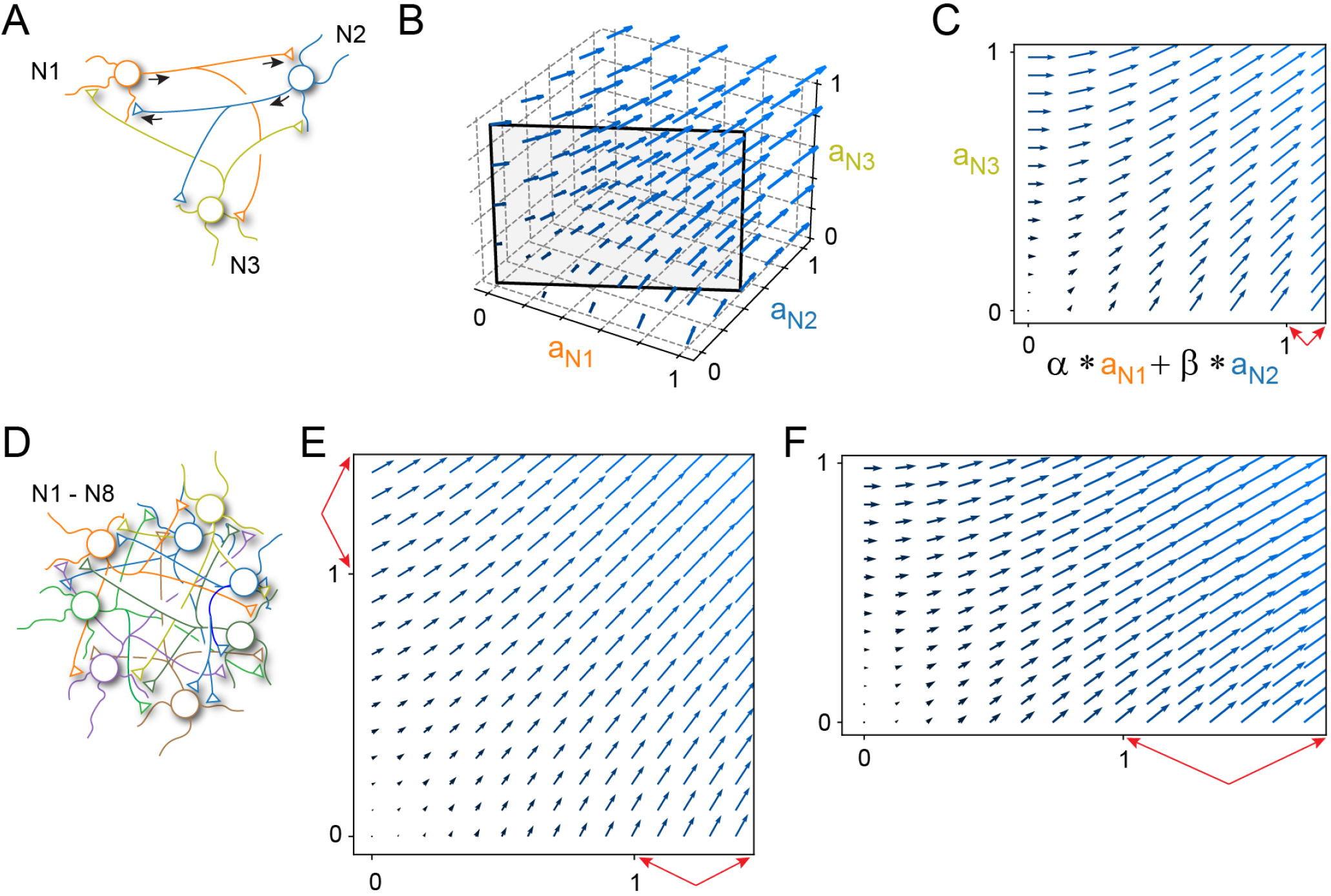
The axes of extracted planes can represent the activity of more than one neuron at a time. (A) The 3-dimensional excitatory network. (B) Vector field representation of the entire 3-dimensional state space with a plane which holds information about the activity of two neurons in one of its dimensions in its x-axis. (C) The 2D plane extracted from the vector field. The x-axis is a linear combination of two out of the three neurons as indicated (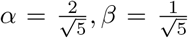). The red arrows indicate that the x-axis as a consequence has a maximal value that exceeds 1. (D) An eight neuron excitatory network. (E) Extracted plane from the 8-dimensional network with two neurons represented on each axis (N1,N2 and N3,N4). (F) Plane extracted from the same network but where one axis represents the combined activity of 3 neurons while the other just a single one (N1-N3 and N4). Across all panels, the synaptic weights were uniformly set to 0.1, all perpendicular neurons had an activity of 0.

### 3.2 Location of the critical point depends on synaptic weights

A critical point is a point in the state space where no change in activity occurs (Figure 1). In visual terms, this means that the vector lengths at that position are zero. Due to the neuron model definition (Equation 1), any change in the synaptic weights or in the activity of the perpendicular neuron(s) can cause the critical point to shift locations.

We can analytically determine the location of the critical point by only taking the numerator of the neuron model (Equation 1) into consideration:

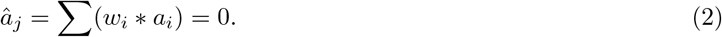

Given a network which is either fully excitatory or fully inhibitory, we can see that the only solutions will be

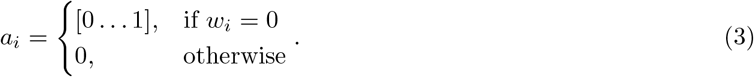

If the synaptic weights are 0, we are left with disconnected neurons. Therefore, in a fully excitatory or in a fully inhibitory network where the neurons are fully connected, the location of the critical point will be at zero (the origin). This motivated us to explore whether it would be possible to displace the critical point from the origin, if the neural network contains both excitatory and inhibitory neurons.

First we consider a 3 neuron network consisting of 2 excitatory neurons and 1 inhibitory neuron (Figure 4A). If we make the inhibitory neuron perpendicular, then the vector field plane describes the interaction between N1 and N2, at a given selected activity level of the inhibitory neuron (N3). The location of the critical point now indeed can be offset from zero activity in the two neurons N1 and N2 (Figure 4B). In this case, when we have outgoing synaptic connections of equal weight (w1=w2) from the inhibitory neuron to the two excitatory neurons, they will be impacted equally. Now, if we increase the weights of the inhibitory neuron (while maintaining the condition w1=w2), the location of the critical point will move only along the central diagonal (Figure 4B) in the neuron activity space of N1 and N2. The surrounding vector field also dramatically changes its structure, as shown by the two examples in Figure 4C.

**Figure 4:**
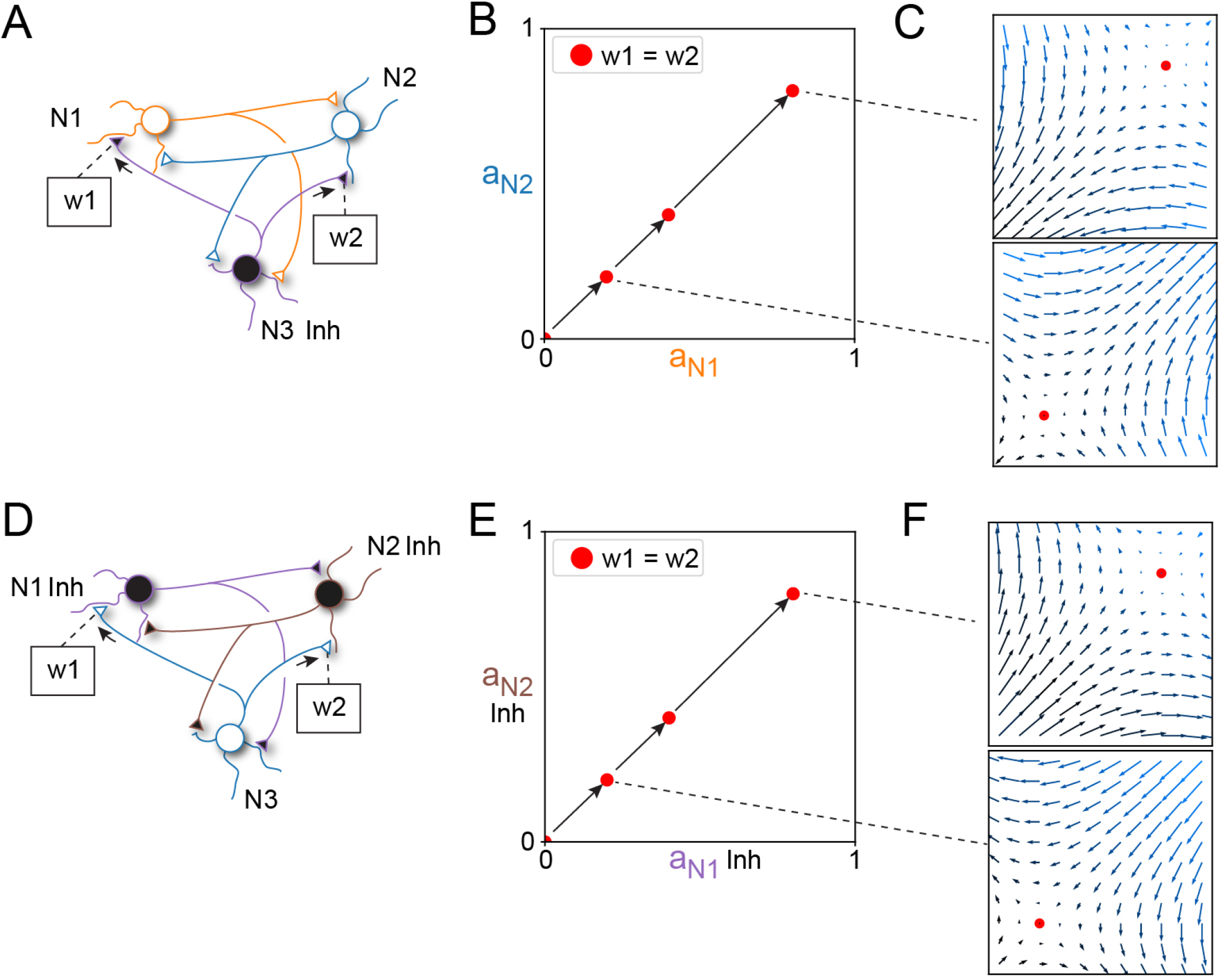
In networks with both excitatory and inhibitory neurons the critical point can be controlled. (A) A three neuron network with one inhibitory neuron (Inh). The weights of the synapses made by the inhibitory neuron are indicated as w1 and w2. (B) Extracted plane with the activity of the two excitatory neurons represented on the axes, and the inhibitory neuron represented on the perpendicular axis. When the weights w1 and w2 were increased from zero (both weights were always equal), the location of the critical point moved upwards along the central diagonal of the vector field (individual vectors are not shown for clarity). For the three illustrated points, the inhibitory weights were –0.2, –0.4 and –0.8. (C) Two examples of the full vector fields at different weight magnitudes of w1 and w2. (D) A network of two inhibitory neurons and one excitatory neuron, where w1 and w2 were instead the weights of the excitatory synapses. (E) Similar effects as in B arise when the excitatory weights of the perpendicular neuron are increased. The synaptic weight magnitudes of the perpendicular neuron are the same as in B. (F) Examples of full vector fields. Across all panels the weights between N1 and N2 and the activity of the perpendicular neuron were fixed at 1.

We can also flip the tables, and instead having two inhibitory neurons as N1 and N2, and the perpendicular neuron (N3) now being excitatory, with equal weights (w1=w2) on N1 and N2 (Figure 4D-F). Increasing the excitatory weights (while maintaining w1=w2) has the same effect on the critical point as earlier, i.e. its location move upwards along the central diagonal in the vector field (Figure 4B,E). Importantly, however, the structures of the vector fields, again provided as two examples, are ‘inverted’ for the inhibitory network (see the directions of the vectors; Figure 4C,F).

If the synaptic weights of the perpendicular neuron are not uniform, the critical point location will be displaced from the central diagonal because of the greater impact the perpendicular neuron has on one of the neurons. The magnitude of the displacement from the diagonal depends on the relative size of the outgoing synaptic weights: the larger the difference between those weights (the skewness), the larger the displacement from the central diagonal (Figure 5A,B). As a general rule, the skewness of the outgoing synaptic weights of the perpendicular neuron on the neurons represented on the respective axes is what determines the extent of displacement from the central diagonal.

**Figure 5:**
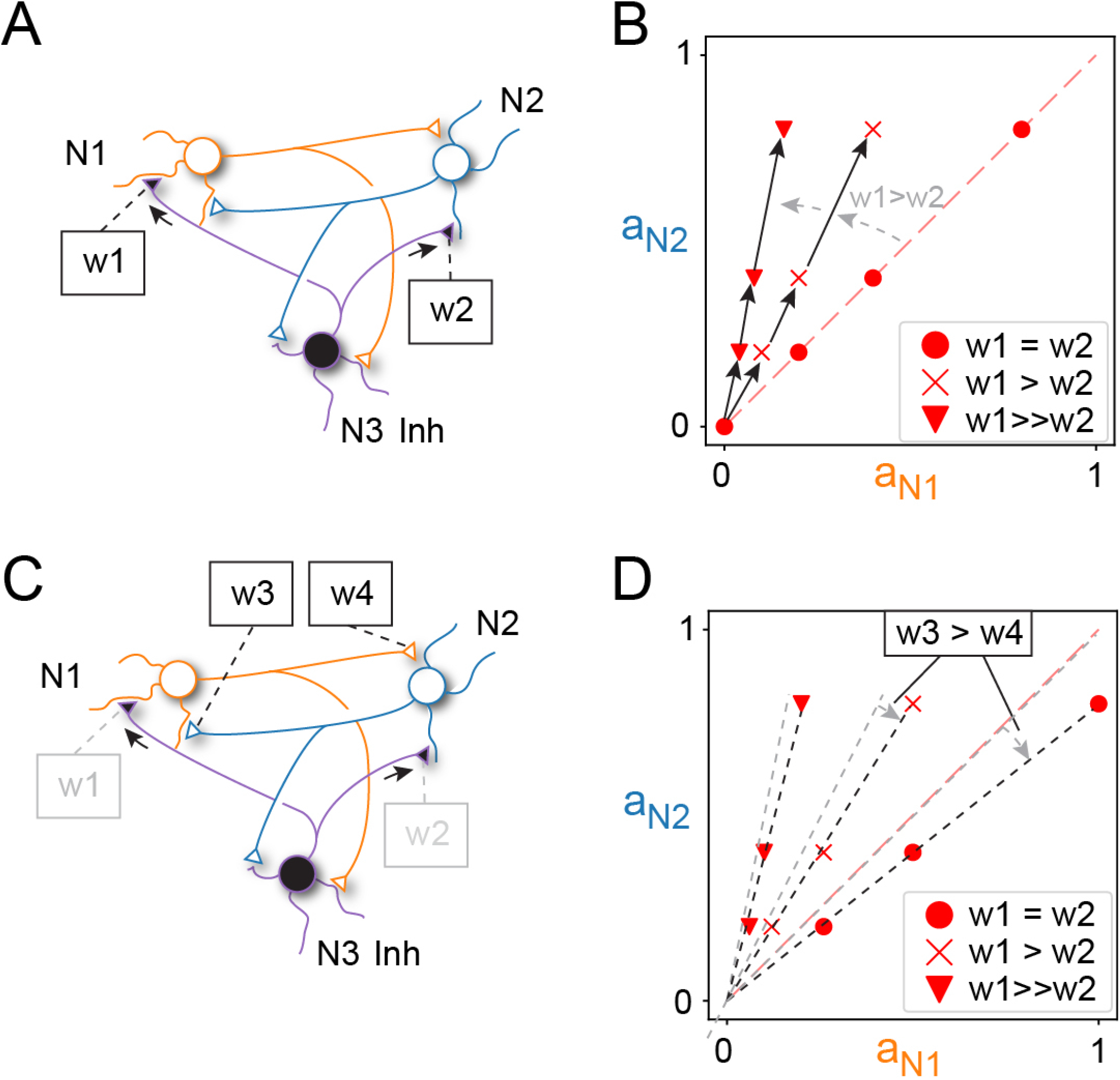
The location of the critical point can shift dramatically when the weights are non-uniform. (A) A network with two excitatory and one inhibitory neuron. Synaptic connections between N1 and N2 were uniformly set to 1. (B) The location of the critical point is offset from the central diagonal when the synaptic weights w1 and w2, formed by the perpendicular neuron, are skewed rather than equal. Each symbol indicates a skewness level, a series of a symbol indicates the location of the critical point when the weights of w1 and w2 were increased proportionally. The weight magnitudes of w1 were set to 0.2, 0.4, 0.8; w2 was scaled down with the following proportions: 1 (unchanged), 2, 5. (C) The same network indicating the synaptic weights w3 and w4 for the synapses made between the non-perpendicular neurons. (D) Impact of making the weights w3 and w4 non-uniform (w3=1, w4=0.8). The arrows indicate the impact on the locations of the critical points when the weights were made non-uniform for the same skewness levels of w1 and w2 as in B. Across all panels the activity of the perpendicular neuron were fixed at 1.

However, the interactions are also influenced by the weights between the non-perpendicular neurons (the neurons represented on the axes). Previously, these weights were equal (w3 = w4) (Figure 5C). If we instead skew these weights, the movement of the critical point will deviate from its original diagonal path observed when the weights w1 and w2 were proportionally increased (Figure 5D). Thus, the trajectory of the critical point, when varying the weights of the perpendicular neuron (w1, w2), is also affected by the weight distribution between the two non-perpendicular neurons. In our 3-dimensional network example, the general relationship between the critical point location, in terms of neuron activity levels, and incoming synaptic weights can be expressed as

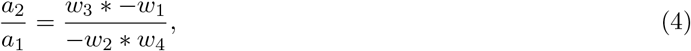

where *a*_1_, *a*_2_ are the activity levels of N1 and N2, the synaptic weights made by the perpendicular neuron N3 are *w*_1_, *w*_2_, while *w*_3_, *w*_4_ are the weights between N1 and N2 (as shown in Figure 5C).

### 3.3 Location of the critical point can be controlled by the activity level

While it is the synaptic weights that govern the structure of the vector field, the specific activity levels of the neurons determine which subspace of the vector field that the network is located in. Through manipulation of the perpendicular neuron activities, we can modify the position of the critical point and thus the structure of the underlying vector field within the extracted plane. The vector field structure is what impacts the evolution of the population level activity (i.e. the network activity state) in that plane, within the constraints that the synaptic weight distributions would allow for. This is illustrated in Figure 6, where we used a 4 neuron network with two perpendicular inhibitory neurons. The activities of the two perpendicular neurons were altered independently of each other to obtain different combinations of activity (Figure 6B). As shown in Figure 6C, the location of the critical point now travelled over a very large range of the activity space of the two non-perpendicular neurons, and the structure of the surrounding vector field also changed dramatically (Figure 6D).

**Figure 6:**
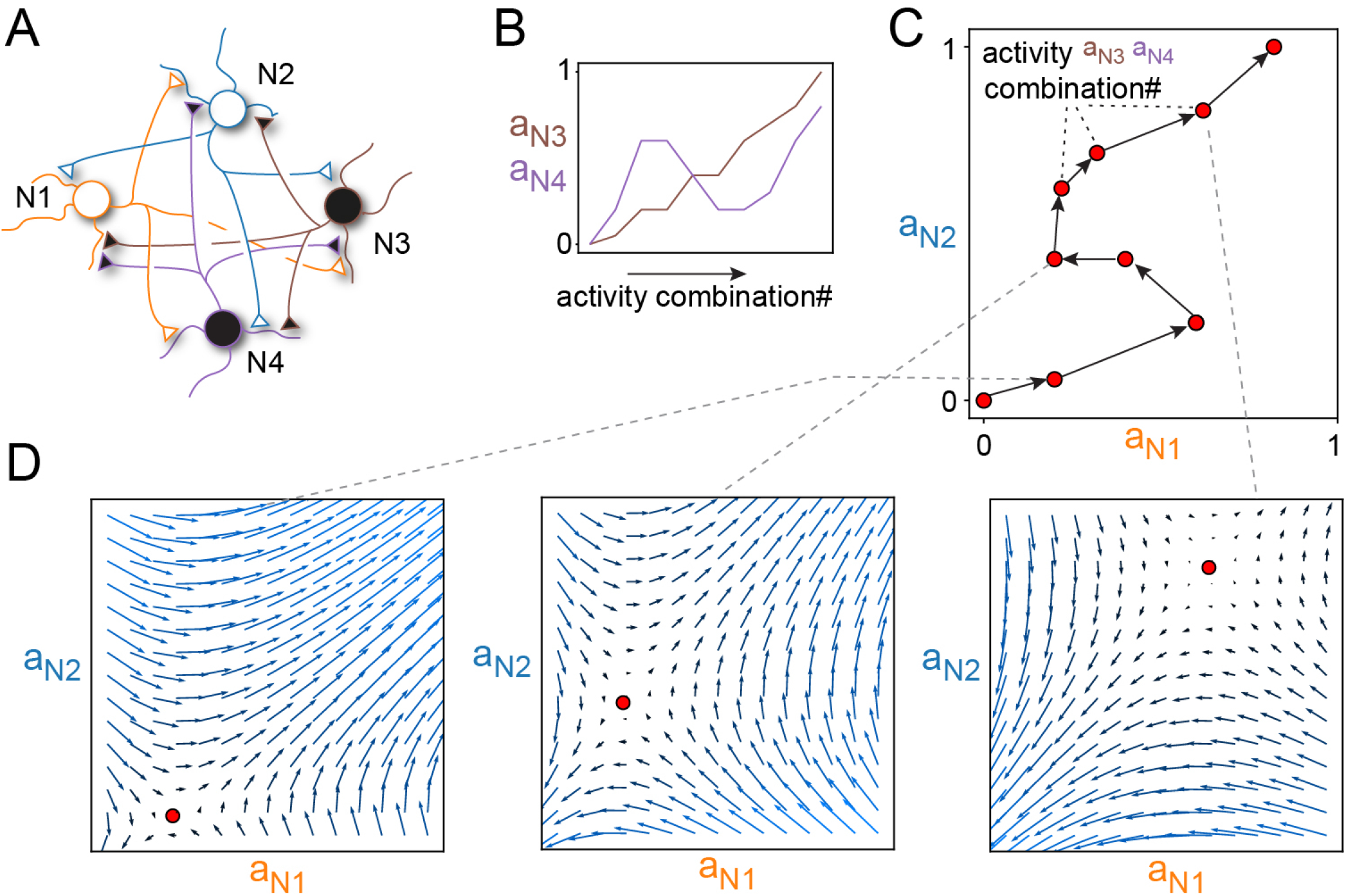
The activity levels of the perpendicular neurons dynamically control the critical point location. (A) Four-dimensional network consisting of two excitatory and two inhibitory neurons. All synaptic weights were set to 0.5, except the synaptic weights from N3 to N2 and N4 to N1 which were reduced to 0.01. (B) Predefined activity level trajectories of the two inhibitory neurons being perpendicular to the extracted plane. (C) Critical point trajectory within the selected plane resulting from the activity level settings in B. (D) Examples of full vector fields from three of the locations of the critical point.

## 4 Discussion

Introducing a conceptual framework to analyze neuron population level interactions within recursive networks, we showed that an inevitable emergent effect of the synaptic weights and the input-output functions of the neurons are vector fields, which define in which direction the activity state of the whole network will be pushed. In other words, the vector at any given point in the network state space will impact how the activity distribution across the neuron population (i.e. the network state) evolves over time. Since basically all aspects of what we consider brain behavioral function depends on controlling its spatiotemporal patterns of neuron activity, we believe that this conceptual framework can be a useful starting point to analyze the properties of high-dimensional networks. The underlying representation obtained with this approach is the complete, high-dimensional vector field of the network, and it can thus help avoiding discarding crucial information regarding its behavior [3], which is a known problem with the dimensionality reduction methods used in the analysis of multi-neuron interactions.

In order to get a more detailed understanding of the network state space and its dependence on connectivity and neuron activity, it is important to be able to visualize the vector field. However, as visualizing in high dimensions is in principle impossible, we introduced a planar subspace extraction tool for visualizing the combined action of a large number of neurons. This plane extraction tool can be used to illustrate any plane within the high dimensional network activity space. The extracted plane represents the interactions between two groups of neurons (one per axis), where a group could be a single neuron or, alternatively, any linear combination of neurons in the network (Figure 3). The choice of neuron groups is what defines the orientation of the extracted plane within the high-dimensional network activity space. With this tool we can sample an arbitrary number of diverse planes from the complete high-dimensional vector field to obtain an overview of the underlying mechanisms that governs the activity distribution across the entire neuron population.

Under certain circumstances, a critical point is present within the vector field of the extracted plane (Figure 2). Given a fully connected, excitatory network, the critical point always ends up in the origin (Figure 1) regardless of network dimensionality (unless any of the neurons only have non-zero activity within the visualized plane, as in Figures 2,3). Beyond that point, all vectors point in the direction of endlessly increasing excitation (saturation), which in biological systems would lead to an epileptic state, a natural consequence of the positive feedback loops created in recursive excitatory networks. Similarly, in a fully inhibitory network the critical point is also located to the origin, but all neuron activity is pushed towards zero, which of course would correspond to an inactive brain. If the network instead contains both excitatory and inhibitory neurons, we showed that it is possible to displace the critical point from the origin (Figure 4) and to reshape the structure of the vector field around it. In this way, we can avoid obtaining networks which will inevitably be driven towards either zero neuron activity or towards saturation. Moreover, any change in neuron activity will now impact the location of the critical point and the structure of the vector field (Figure 6). Since the impact of the vector field is to push the network towards new activity states, thereby inevitably also changing the critical point, it should be possible to design such excitatory-inhibitory networks where the activity of the network constantly changes into new patterns, as in the brain *in vivo* [9, 17, 8, 18], as a consequence of that the network would constantly be ‘chasing its own tail’ (i.e. the critical point).

A wider range of variations of the critical point location, and its surrounding vector field, could be obtained by having some large and some small synaptic weights (perhaps even zero weight), or in other words, skewing the network weights (Figures 4, 5). This means that networks with skewed weights, rather than uniform weights, can reach a wider range of their activity state space. This would in turn translate to that these networks would have a greater representational capacity.

An implication of the vector field analogy is that in the brain, the constantly evolving activity distribution across the neuron population will be a consequence of the synaptic connectivity landscape of the network. These ‘trajectories’ through the state space are the solutions provided by the network to given sensory inputs, at the cortical level [8, 9] as well as in the spinal cord circuitry [11, 13, 12]. These network solutions could essentially be the primary basis for how behavioral decisions are made. They will control how the activity distributions of the neuron population will evolve. The activity distributions in turn translate to for example in what order and with what magnitude we activate specific muscles. Muscle activation patterns are an expression of behavioral choices so this is a concrete example of the direct relevance of the vector fields to understand behavior. But the vector field representation can also explain perceptual function where a sensory effect is immediately registered by the network but may be saved to impact behavioral decisions at a later time point [19].

## 5 Conclusion

We used a simple neuron model to investigate effects that would inevitably emerge from interactions in neuron populations. We found that the distribution of synaptic weights determines the structure of a vector field, while the activity distribution across the neuron population –i.e., the network’s position within its activity state space— determines a vector which will influence network activity, thereby shaping how the activity distribution across the network evolves. These principles apply also to real brain networks, but there we have different factors that can introduce additional complexity. For instance, synaptic weights between neurons must be known or estimated, and the input-output functions may vary across neuron types. Some neurons may exhibit non-linear activation functions, and others may communicate via metabotropic rather than ionotropic synapses, which have lower instantaneous effects but influence the target neuron over a longer duration and can alter the gain of their activation function [20]. Regardless of these complexities, our results indicate that synaptic weight distributions are the primary factor driving the evolution of neuronal activity distribution – which in turn is likely to be one of the most critical elements in defining brain function.

## References

[1] Mehrdad Jazayeri and Srdjan Ostojic. Interpreting neural computations by examining intrinsic and embedding dimensionality of neural activity. Current opinion in neurobiology, 70:113–120, 2021.

[2] Mikail Khona and Ila R Fiete. Attractor and integrator networks in the brain. Nature Reviews Neuroscience, 23(12):744–766, 2022.

[3] Sofie S Kristensen, Kaan Kesgin, and Henrik Jörntell. High-dimensional cortical signals reveal rich bimodal and working memory-like representations among s1 neuron populations. Communications biology, in press, 2024.

[4] Rodney Douglas, Christof Koch, Misha Mahowald, and Kevan Martin. The role of recurrent excitation in neocortical circuits. In Models of cortical circuits, pages 251–282. Springer, 1999.

[5] Tom Binzegger, Rodney J Douglas, and Kevan AC Martin. A quantitative map of the circuit of cat primary visual cortex. Journal of Neuroscience, 24(39):8441–8453, 2004.

[6] Nir Kalisman, Gilad Silberberg, and Henry Markram. The neocortical microcircuit as a tabula rasa. Proceedings of the National Academy of Sciences, 102(3):880–885, 2005.

[7] Jonas MD Enander, Anton Spanne, Alberto Mazzoni, Fredrik Bengtsson, Calogero Maria Oddo, and Henrik Jörntell. Ubiquitous neocortical decoding of tactile input patterns. Frontiers in cellular neuroscience, 13:140, 2019.

[8] Leila Etemadi, Jonas MD Enander, and Henrik Jörntell. Remote cortical perturbation dynamically changes the network solutions to given tactile inputs in neocortical neurons. Iscience, 25(1), 2022.

[9] Johanna Norrlid, Jonas MD Enander, Hannes Mogensen, and Henrik Jörntell. Multi-structure cortical states deduced from intracellular representations of fixed tactile input patterns. Frontiers in cellular neuroscience, 15:677568, 2021.

[10] Anders Wahlbom, Jonas MD Enander, Fredrik Bengtsson, and Henrik Jörntell. Focal neocortical lesions impair distant neuronal information processing. The Journal of physiology, 597(16):4357–4371, 2019.

[11] Matthias Kohler, Fredrik Bengtsson, Philipp Stratmann, Florian Röhrbein, Alois Knoll, Alin Albu-Schäffer, and Henrik Jörntell. Diversified physiological sensory input connectivity questions the existence of distinct classes of spinal interneurons. Iscience, 25(4), 2022.

[12] Marco Santello, Gabriel Baud-Bovy, and Henrik Jörntell. Neural bases of hand synergies. Frontiers in computational neuroscience, 7:23, 2013.

[13] Matthias Kohler, Florian Röhrbein, Alois Knoll, Alin Albu-Schäffer, and Henrik Jörntell. The bcm rule allows a spinal cord model to learn rhythmic movements. Biological Cybernetics, 117(4):275–284, 2023.

[14] Udaya B Rongala, Jonas MD Enander, Matthias Kohler, Gerald E Loeb, and Henrik Jörntell. A non-spiking neuron model with dynamic leak to avoid instability in recurrent networks. Frontiers in computational neuroscience, 15:656401, 2021.

[15] Ingrid HCHM Philippens. Marmosets in neurologic disease research: Parkinson’s disease. In The Common Marmoset in Captivity and Biomedical Research, pages 415–435. Elsevier, 2019.

[16] Ivan Pavlov, Leonid P Savtchenko, Dimitri M Kullmann, Alexey Semyanov, and Matthew C Walker. Outwardly rectifying tonically active gabaa receptors in pyramidal cells modulate neuronal offset, not gain. Journal of Neuroscience, 29(48):15341–15350, 2009.

[17] Carsen Stringer, Marius Pachitariu, Nicholas Steinmetz, Charu Bai Reddy, Matteo Carandini, and Kenneth D Harris. Spontaneous behaviors drive multidimensional, brainwide activity. Science, 364(6437):eaav7893, 2019.

[18] Nghia D Nguyen, Andrew Lutas, Oren Amsalem, Jesseba Fernando, Andy Young-Eon Ahn, Richard Hakim, Josselyn Vergara, Justin McMahon, Jordane Dimidschstein, Bernardo L Sabatini, et al. Cortical reactivations predict future sensory responses. Nature, 625(7993):110–118, 2024.

[19] Andy Clark. Whatever next? predictive brains, situated agents, and the future of cognitive science. Behavioral and brain sciences, 36(3):181–204, 2013.

[20] Philipp Stratmann, Alin Albu-Schäffer, and Henrik Jörntell. Scaling our world view: how monoamines can put context into brain circuitry. Frontiers in cellular neuroscience, 12:506, 2018.

